# Informing Extinction Risk: Summarizing Population Viability through a Meta-analysis of Multiple Long-term Monitoring Programs for a Declining Estuarine Fish Species

**DOI:** 10.1101/2023.04.17.537200

**Authors:** Vanessa D. Tobias, Ernest Chen, James Hobbs, Michael Eakin, Steven Detwiler

## Abstract

Decisions about whether to designate a species for conservation and protection depend on the ability to summarize their population trajectories and their risk of extinction. Such decisions may rely on quantitative population viability analyses based on a time series of abundance index values that are derived from a monitoring program. In the case of the Longfin Smelt (*Spirinchus thaleichthys*), a decision to protect a distinct population segment of the species under the U.S. Endangered Species Act was informed by several indices of population abundance. In this paper, we combined individual population viability analyses into a single metric for extinction risk using a meta-analysis framework. Individual monitoring surveys for this species generally agreed that the trajectory of abundance was downward and combining data from all of the surveys produced a better summary of the population growth rate. We also used the population growth rates in a simulation to estimate the probability that the abundance of Longfin Smelt dropped too low to recover. We found that this probability of quasi-extinction was substantial, exceeding 20% over two decades. This study demonstrates a practical way that having multiple sources of information creates better information about the trajectory of a population. Individually, the surveys contribute information about specific life stages or ages to our understanding of the population. Combined into one metric and an associated graphical summary, this analysis succinctly communicates risk and creates a benchmark for evaluating future management decisions.

## Introduction

Designating species as endangered or threatened may serve to highlight the need for conservation in a scientific, popular, or political context, as is the goal of the IUCN Red List (Mace et al. 2008), or to confer regulatory protections, as is the case for the U.S. Endangered Species Act of 1973 (ESA) and the California Endangered Species Act of 1970 (CESA). For the ESA, the decision to include a species under the protection of the ESA is made by the Secretaries of the Interior and Commerce, acting through the U.S. Fish and Wildlife Service (USFWS) and National Oceanic and Atmospheric Administration, based on an assessment of the best available scientific and commercial data for the species (United States 1983; Smith et al. 2018). The USFWS recently implemented a new framework for decision making related to the ESA. Under the new approach, experts gather the best available information and data on a species into a Species Status Assessment (SSA) report (Smith et al. 2018). The SSA framework supports decision-making by describing the species needs and life history as well as current and potential future threats to its continued existence in the wild. SSAs also include assessments of the future condition of the species in light of anticipated changes in the magnitude of extant or future stressors. In some cases, this may be a qualitative description of the trajectory of the species, or it could be a quantitative assessment of population viability depending on the quantity and quality of data available (Smith et al. 2018).

Population viability analyses (PVAs) can take many forms, depending on the questions of interest and the data that are available (Morris and Doak 2002), and they have been applied to a wide range of species including insects, fish, reptiles, mammals, and birds (e.g., Shaffer 1983 on grizzly bears, Schultz and Hammond 2003 on butterflies, Tucker et al. 2021 on boa constrictors). Many monitoring programs collect data that is used to assess population status and trends in abundance for species of management concern. From these, a time series of abundance indices can be used to calculate population growth rates and to forecast the risk of extinction into the future through the application of a population viability analysis (PVA). Different methods for conducting PVAs have been developed for different types of available data. A count based PVA is classically applied to census data (counts of an entire population), but it is not necessary to count the entire population. A count based PVA can also be applied to index values, where a population index represents some portion of the total population as long as the proportion of the population that is observed remains relatively constant over time (Morris and Doak 2002). When an SSA includes a quantitative assessment, the experts may need to decide how to incorporate all of the relevant data where multiple long-term datasets exist. In some cases, the textbook PVA methods need to be modified to incorporate data from multiple sources and of differing data types. For example, a combination of available datasets on black rail (*Laterallus jamaicensis*) observations from across their range were combined into a dynamic model of demographic occupancy (McGowan et al. 2020). Population estimates of gopher tortoise (*Gopherus polyphemus*) from multiple sites have been combined into a spatially explicit PVA that is responsive to multiple population threats (Folt et al. 2022). A PVA was also developed that combined habitat area and egg counts in a Bayesian analysis for a population of monarch butterflies (*Danaus plexippus*) that migrates from Mexico to the Midwestern United States (Semmens et al. 2016).

In the fall of 2022, the USFWS completed an SSA for the San Francisco Bay-Delta distinct population segment (DPS) of the Longfin Smelt (*Spirinchus thaleichthys*) and proposed listing it as endangered under the ESA. As is the case for many other species of concern, ecological monitoring provides data on the abundance of Longfin Smelt in the San Francisco Estuary (also known as the San Francisco Bay-Delta). The San Francisco Estuary is one of the most highly studied and monitored ecosystems in the world. Within the Estuary, a science consortium called The Interagency Ecological Program (IEP) conducts a suite of fish monitoring surveys in the San Francisco Estuary, some of which have been collecting data for over 50 years. Several of the IEP long-term monitoring surveys provide indices of Longfin Smelt abundance in the San Francisco Estuary. Some of these indices are calculated and published annually (e.g., the Fall Midwater Trawl [FMWT] abundance index), but another was calculated for this analysis (the 20-mm Survey). An index of abundance need not be a formal index at all; a summary of catch per unit effort (CPUE) can also be thought of as a population index. These long-term monitoring datasets create an opportunity to investigate trends in abundance and to estimate the probability of long-term viability of many species of management concern. Longfin Smelt abundance has declined substantially since monitoring began (Sommer et al. 2007; Rosenfield and Baxter 2007; Nobriga and Rosenfield 2016). Despite consensus that abundance has trended downward, the fluctuations around that trend can make it difficult to quantitatively assess the risk of extinction. For example, it may be difficult to see the effects of one or two “good” years on the overall trajectory of abundance.

In this paper we present a novel analysis of the long-term monitoring data for the San Francisco Bay-Delta DPS of Longfin Smelt which was developed for the SSA. Specifically, we addressed the following questions that are relevant to the SSA: (1) Does the suite of available monitoring data agree on a trajectory for the abundance of Longfin Smelt? (2) What information do we gain by having multiple surveys that track Longfin Smelt abundance? (3) Is the population at risk of extinction within the next two decades? To answer these questions, we summarize the information contained in the monitoring data by applying a count based PVA to the IEP’s monitoring data for Longfin Smelt. Applying the PVA method to several datasets that index the abundance of Longfin Smelt captures the landscape of available information and may be used to make decisions about the status of the species or about management actions. It also presents a way to synthesize the evidence of the direction and magnitude of change in Longfin Smelt abundance from indices calculated using various methods and that exist on varying scales. We further synthesize population growth rates by conducting a meta-analysis on these metrics.

## Methods

### Study Species

Longfin Smelt (*Spirinchus thaleichthys*) is a small (≤150 mm FL), facultatively anadromous, pelagic fish with an up to 3 year lifespan (Baxter 1999). It is native to coastal waters of the eastern Pacific Ocean and bays and estuaries, ranging from Monterey Bay, California, to Alaska (Moyle 2002). Both anadromous and resident populations exist at various locations, and the DPS in the San Francisco Estuary is anadromous (Dryfoos 1965; Moulton 1974; Rosenfield and Baxter 2007). The population of Longfin Smelt in the San Francisco Estuary is genetically distinct from other populations, but there is evidence of outward gene flow from the San Francisco Estuary population to populations in Humboldt Bay and the Columbia River (Garwood 2017; Sağlam et al. 2021). Based in part on the substantial decline in abundance, Longfin Smelt was listed as threatened state-wide under the CESA in 2009 by the California Department of Fish and Game [now the California Department of Fish and Wildlife] ([OAL] Office of Administrative Law 2010), and the USFWS determined that protection of the Bay-Delta DPS of the Longfin Smelt under the ESA was warranted but precluded by higher priority actions (U. S. Office of the Federal Register 2012). The novel analysis reported herein was utilized to help inform the extant and future extinction risk for the DPS and a final determination by USFWS as either threatened or endangered under the ESA.

### Data

Several long-term monitoring surveys produce abundance indices for Longfin Smelt in San Francisco Bay and the Sacramento San Joaquin Delta. We used abundance indices from the FMWT (1 index)(CDFW and IEP 2021a), San Francisco Bay Study (SBFBS: 6 indices) (CDFW and IEP 2021b), and the 20-mm Survey (1 index) (CDFW and IEP 2021c) to estimate apparent annual population growth rates for various life stages of Longfin Smelt (Figure 1). The Bay Study tracks three age classes using two gear types to produce six indices of abundance. The FMWT produces an index that combines all age classes. The 20-mm Survey focuses on juvenile fishes.

**Figure 1:**
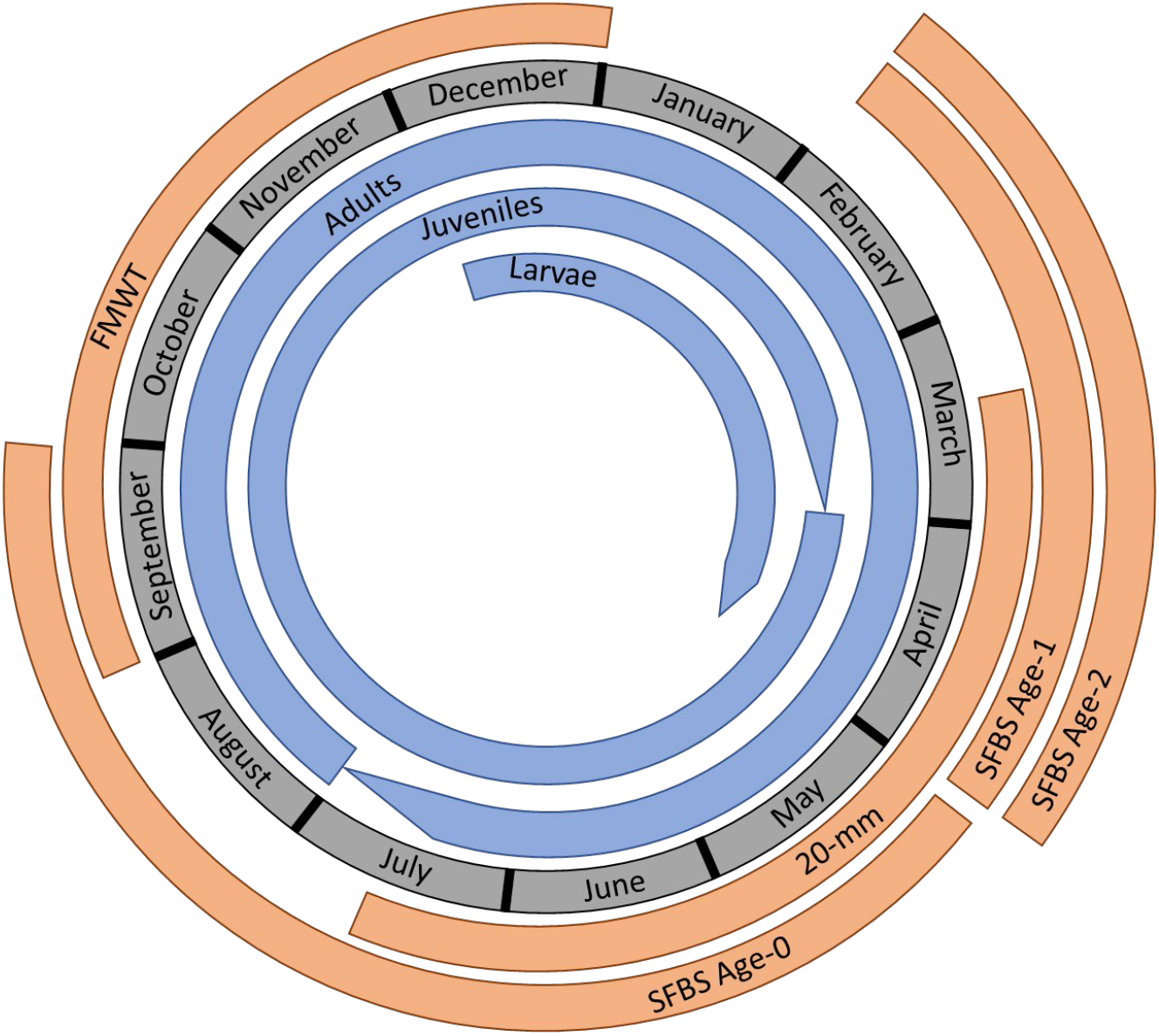
Presence of Longfin Smelt life stages (blue; inside of the ring of months) in the estuary and the timing of monitoring surveys (orange; outside) throughout a calendar year. Bands for life stages taper at the end of the time frame to differentiate the beginning from the end of the bands.

The 20-mm Survey does not produce an index of Longfin Smelt abundance, so we adapted the methods used for the Delta Smelt index to create a 20-mm survey abundance index for Longfin Smelt. We calculated CPUE for index stations (Longfin Smelt catch per 10,000 m^3^ of water), and log transformed the CPUE values, using the usual 20-mm method (log_10_(CPUE + 1)). We calculated the mean of the log-CPUE values for each month within a year and then back-transformed the mean values to put them back on the untransformed density scale. The 20-mm survey index for Delta Smelt bases the selection of surveys on the size of Delta Smelt. For this survey, we used the maximum density value as the index of abundance rather than a mean of specific months. We did this partially for simplicity and partially because using the maximum value reduces the impact of any issues with changing timing of Longfin Smelt presence in the 20-mm survey sampling area.

### Trends in Abundance Indices

We tested for monotonic trends in the individual time series (Table 1) using a Mann-Kendall test (Pohlert 2020). Missing values at the beginning and end of each time series were trimmed; missing values inside of the time series were filled with the mean of the two closest non-missing values (i.e., usually the prior and subsequent year unless there were consecutive missing values).

**Table 1:**
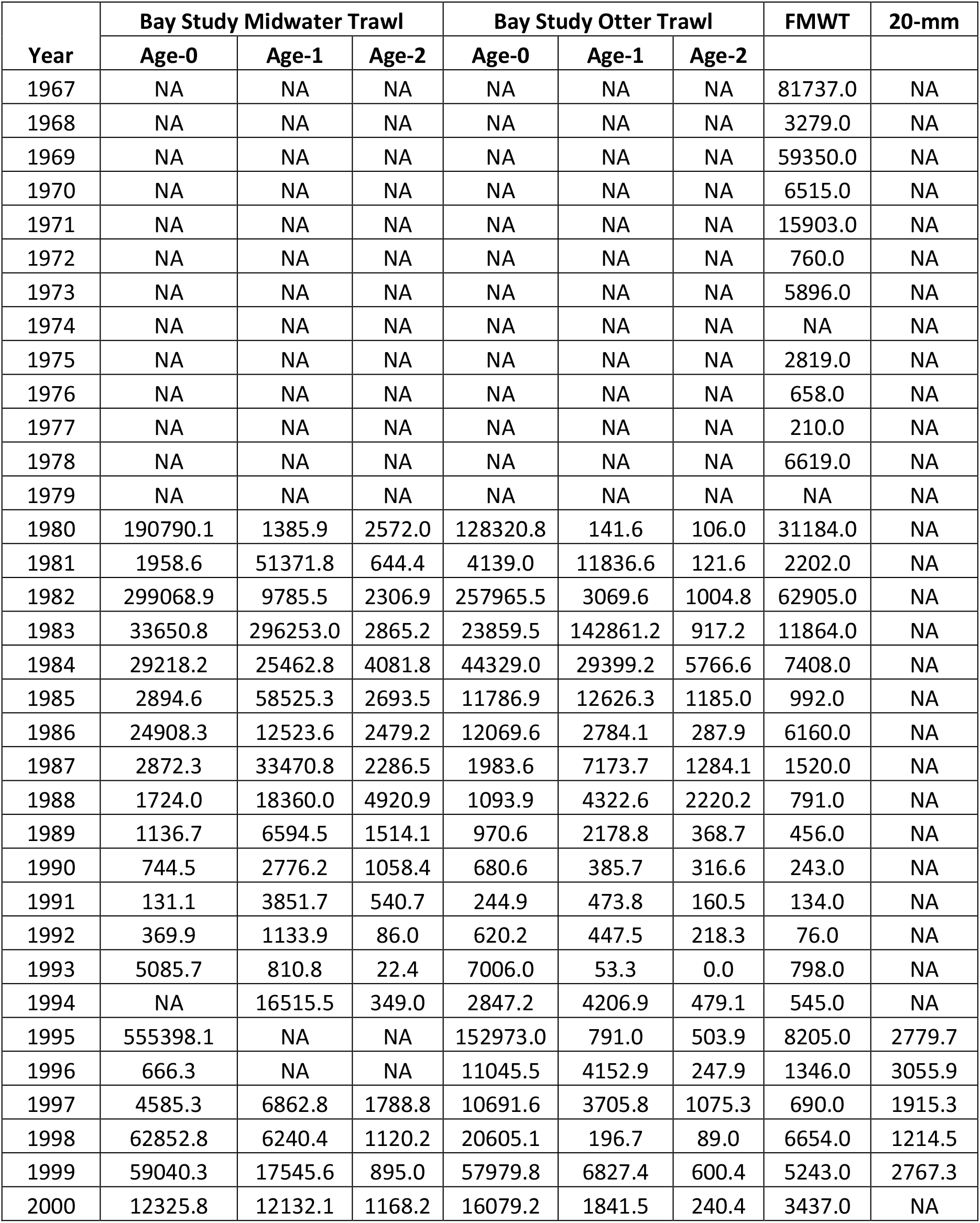
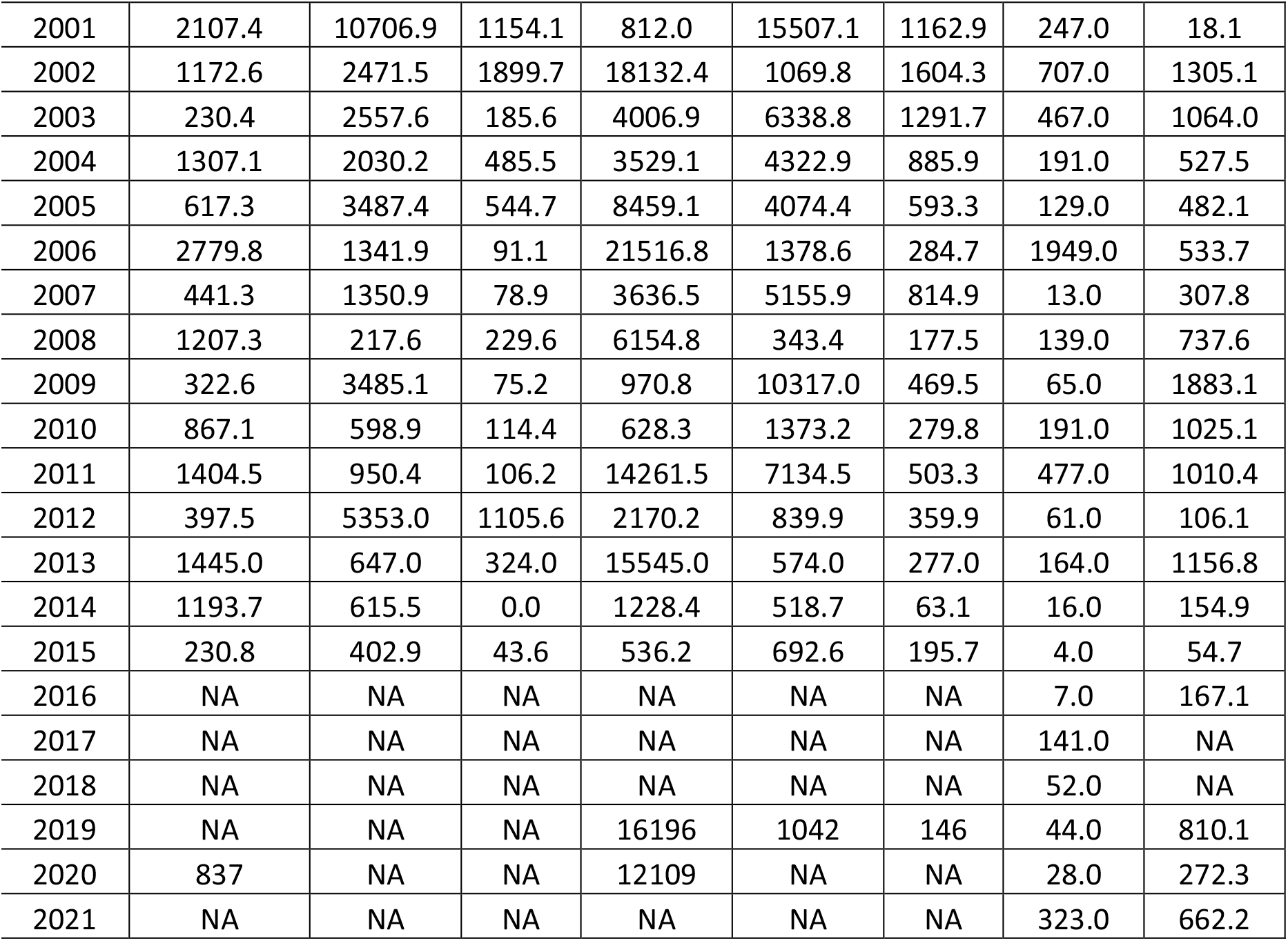
Abundance index data that were used in the population viability analysis. Years when indices were not calculated are denoted by NA: Bay Study began surveying in 1980; 20-mm began sampling in 1995; a combination of sampling gear issues and the COVID-19 pandemic prevented calculation of many abundance indices from 2016-2021.

| Year | Bay Study Midwater Trawl |  |  | Bay Study Otter Trawl |  |  | FMWT | 20-mm |
| --- | --- | --- | --- | --- | --- | --- | --- | --- |
|  | Age-0 | Age-1 | Age-2 | Age-0 | Age-1 | Age-2 |  |  |
| 1967 | NA | NA | NA | NA | NA | NA | 81737.0 | NA |
| 1968 | NA | NA | NA | NA | NA | NA | 3279.0 | NA |
| 1969 | NA | NA | NA | NA | NA | NA | 59350.0 | NA |
| 1970 | NA | NA | NA | NA | NA | NA | 6515.0 | NA |
| 1971 | NA | NA | NA | NA | NA | NA | 15903.0 | NA |
| 1972 | NA | NA | NA | NA | NA | NA | 760.0 | NA |
| 1973 | NA | NA | NA | NA | NA | NA | 5896.0 | NA |
| 1974 | NA | NA | NA | NA | NA | NA | NA | NA |
| 1975 | NA | NA | NA | NA | NA | NA | 2819.0 | NA |
| 1976 | NA | NA | NA | NA | NA | NA | 658.0 | NA |
| 1977 | NA | NA | NA | NA | NA | NA | 210.0 | NA |
| 1978 | NA | NA | NA | NA | NA | NA | 6619.0 | NA |
| 1979 | NA | NA | NA | NA | NA | NA | NA | NA |
| 1980 | 190790.1 | 1385.9 | 2572.0 | 128320.8 | 141.6 | 106.0 | 31184.0 | NA |
| 1981 | 1958.6 | 51371.8 | 644.4 | 4139.0 | 11836.6 | 121.6 | 2202.0 | NA |
| 1982 | 299068.9 | 9785.5 | 2306.9 | 257965.5 | 3069.6 | 1004.8 | 62905.0 | NA |
| 1983 | 33650.8 | 296253.0 | 2865.2 | 23859.5 | 142861.2 | 917.2 | 11864.0 | NA |
| 1984 | 29218.2 | 25462.8 | 4081.8 | 44329.0 | 29399.2 | 5766.6 | 7408.0 | NA |
| 1985 | 2894.6 | 58525.3 | 2693.5 | 11786.9 | 12626.3 | 1185.0 | 992.0 | NA |
| 1986 | 24908.3 | 12523.6 | 2479.2 | 12069.6 | 2784.1 | 287.9 | 6160.0 | NA |
| 1987 | 2872.3 | 33470.8 | 2286.5 | 1983.6 | 7173.7 | 1284.1 | 1520.0 | NA |
| 1988 | 1724.0 | 18360.0 | 4920.9 | 1093.9 | 4322.6 | 2220.2 | 791.0 | NA |
| 1989 | 1136.7 | 6594.5 | 1514.1 | 970.6 | 2178.8 | 368.7 | 456.0 | NA |
| 1990 | 744.5 | 2776.2 | 1058.4 | 680.6 | 385.7 | 316.6 | 243.0 | NA |
| 1991 | 131.1 | 3851.7 | 540.7 | 244.9 | 473.8 | 160.5 | 134.0 | NA |
| 1992 | 369.9 | 1133.9 | 86.0 | 620.2 | 447.5 | 218.3 | 76.0 | NA |
| 1993 | 5085.7 | 810.8 | 22.4 | 7006.0 | 53.3 | 0.0 | 798.0 | NA |
| 1994 | NA | 16515.5 | 349.0 | 2847.2 | 4206.9 | 479.1 | 545.0 | NA |
| 1995 | 555398.1 | NA | NA | 152973.0 | 791.0 | 503.9 | 8205.0 | 2779.7 |
| 1996 | 666.3 | NA | NA | 11045.5 | 4152.9 | 247.9 | 1346.0 | 3055.9 |
| 1997 | 4585.3 | 6862.8 | 1788.8 | 10691.6 | 3705.8 | 1075.3 | 690.0 | 1915.3 |
| 1998 | 62852.8 | 6240.4 | 1120.2 | 20605.1 | 196.7 | 89.0 | 6654.0 | 1214.5 |
| 1999 | 59040.3 | 17545.6 | 895.0 | 57979.8 | 6827.4 | 600.4 | 5243.0 | 2767.3 |
| 2000 | 12325.8 | 12132.1 | 1168.2 | 16079.2 | 1841.5 | 240.4 | 3437.0 | NA |
| 2001 | 2107.4 | 10706.9 | 1154.1 | 812.0 | 15507.1 | 1162.9 | 247.0 | 18.1 |
| 2002 | 1172.6 | 2471.5 | 1899.7 | 18132.4 | 1069.8 | 1604.3 | 707.0 | 1305.1 |
| 2003 | 230.4 | 2557.6 | 185.6 | 4006.9 | 6338.8 | 1291.7 | 467.0 | 1064.0 |
| 2004 | 1307.1 | 2030.2 | 485.5 | 3529.1 | 4322.9 | 885.9 | 191.0 | 527.5 |
| 2005 | 617.3 | 3487.4 | 544.7 | 8459.1 | 4074.4 | 593.3 | 129.0 | 482.1 |
| 2006 | 2779.8 | 1341.9 | 91.1 | 21516.8 | 1378.6 | 284.7 | 1949.0 | 533.7 |
| 2007 | 441.3 | 1350.9 | 78.9 | 3636.5 | 5155.9 | 814.9 | 13.0 | 307.8 |
| 2008 | 1207.3 | 217.6 | 229.6 | 6154.8 | 343.4 | 177.5 | 139.0 | 737.6 |
| 2009 | 322.6 | 3485.1 | 75.2 | 970.8 | 10317.0 | 469.5 | 65.0 | 1883.1 |
| 2010 | 867.1 | 598.9 | 114.4 | 628.3 | 1373.2 | 279.8 | 191.0 | 1025.1 |
| 2011 | 1404.5 | 950.4 | 106.2 | 14261.5 | 7134.5 | 503.3 | 477.0 | 1010.4 |
| 2012 | 397.5 | 5353.0 | 1105.6 | 2170.2 | 839.9 | 359.9 | 61.0 | 106.1 |
| 2013 | 1445.0 | 647.0 | 324.0 | 15545.0 | 574.0 | 277.0 | 164.0 | 1156.8 |
| 2014 | 1193.7 | 615.5 | 0.0 | 1228.4 | 518.7 | 63.1 | 16.0 | 154.9 |
| 2015 | 230.8 | 402.9 | 43.6 | 536.2 | 692.6 | 195.7 | 4.0 | 54.7 |
| 2016 | NA | NA | NA | NA | NA | NA | 7.0 | 167.1 |
| 2017 | NA | NA | NA | NA | NA | NA | 141.0 | NA |
| 2018 | NA | NA | NA | NA | NA | NA | 52.0 | NA |
| 2019 | NA | NA | NA | 16196 | 1042 | 146 | 44.0 | 810.1 |
| 2020 | 837 | NA | NA | 12109 | NA | NA | 28.0 | 272.3 |
| 2021 | NA | NA | NA | NA | NA | NA | 323.0 | 662.2 |

### Population Growth Rates

Annual population growth rates (lambdas) estimates are based on a simple unstructured population time series, where a single estimate of population size (or the size of a segment of the population) is produced annually. We used an exponential growth state space (EGSS) model similar to the REML model which (Staples et al. 2004) proposed as a solution to accounting for observation error in estimating population growth rates, and which they found to produce unbiased estimates of λ.

We used program R (R Core Team 2019) and modified code for an EGSS model from that published by (Humbert et al. 2009) to estimate μ = log(λ) and its associated variance. This method does not allow for zeros in the timeseries so the few instances where the abundance index was zero were replaced with a one. Our approach to calculating population growth rates for multiple surveys was similar to that of Schultz and Hammond (2003), but we took an additional step to summarize the growth rates. We first applied the EGSS model to the time series from each survey. We then conducted a meta-analysis using the mean and variance of population growth rates from the individual surveys as independent measurements of population trajectory and variability. A meta-analysis calculated a pooled estimate of the mean growth rate from results of separate studies. We used the metamean function from the meta package in R (Balduzzi et al. 2019).

### Evaluation of Assumptions

Count PVAs assume that the mean and variance of the population growth rate do not change over time. We assessed this assumption by plotting the lambda values over time. We also checked that the mean population growth rate is density independent. To do this, we graphed the population growth rates against the population size for each year in the datasets and looked for a trend. Graphs for most of the surveys did not show trends, but a couple of plots were inconclusive, so we compared models for density dependence (Beverton-Holt and Ricker) against a density-independent (linear) model for the population growth rate. In order to guard against any undetected density dependence, we followed the advice of Morris and Doak (2002) to first, limit our projections of extinction risk to a relatively short time frame to limit the influence of negative density dependence (i.e., predicting extremely high populations that exceed carrying capacity) and second, set a quasi-extinction threshold to limit the influence of positive density dependence (i.e., Allee effects). To test for environmental autocorrelation, i.e., a tendency for adjacent population growth rates to be more similar to one another, in the timeseries we performed a Durbin-Watson test. When the Durbin-Watson test indicated significant autocorrelation, we used ACF to assess the degree.

### Selection of Quasi-Extinction Thresholds

A quasi-extinction threshold is a population size below which a population cannot recover or when the abundance is so low that plausible management strategies would need to drastically change for the species to recover; for example changing the focus of restoration efforts from habitat management to captive breeding (Tucker et al. 2021). Quasi-extinction is used as the threshold for PVAs rather than full extirpation because this strategy minimizes the uncertainty around population dynamics for small populations (Busch et al. 2013) and because it is consistent with the endangered designation used for the ESA (Boyd et al. 2017). Our analysis uses distinct values for the quasi-extinction thresholds for each of the surveys because the scale of the abundance indices differs across surveys. The following procedure was developed as an attempt to account for the different magnitudes of index values across the surveys. First, we calculated the mean value of all available index values for each survey over 2009-2018. We then multiplied that mean value by 0.01 to represent a value of a major decline in abundance from the recent average. The major decline value was then rounded to zero decimal places to create an integer value. We also set the minimum value to 1 *a priori*, but none of the calculated threshold values was less than 1. Final quasi-extinction threshold values are reported in Table 2.

**Table 2:**
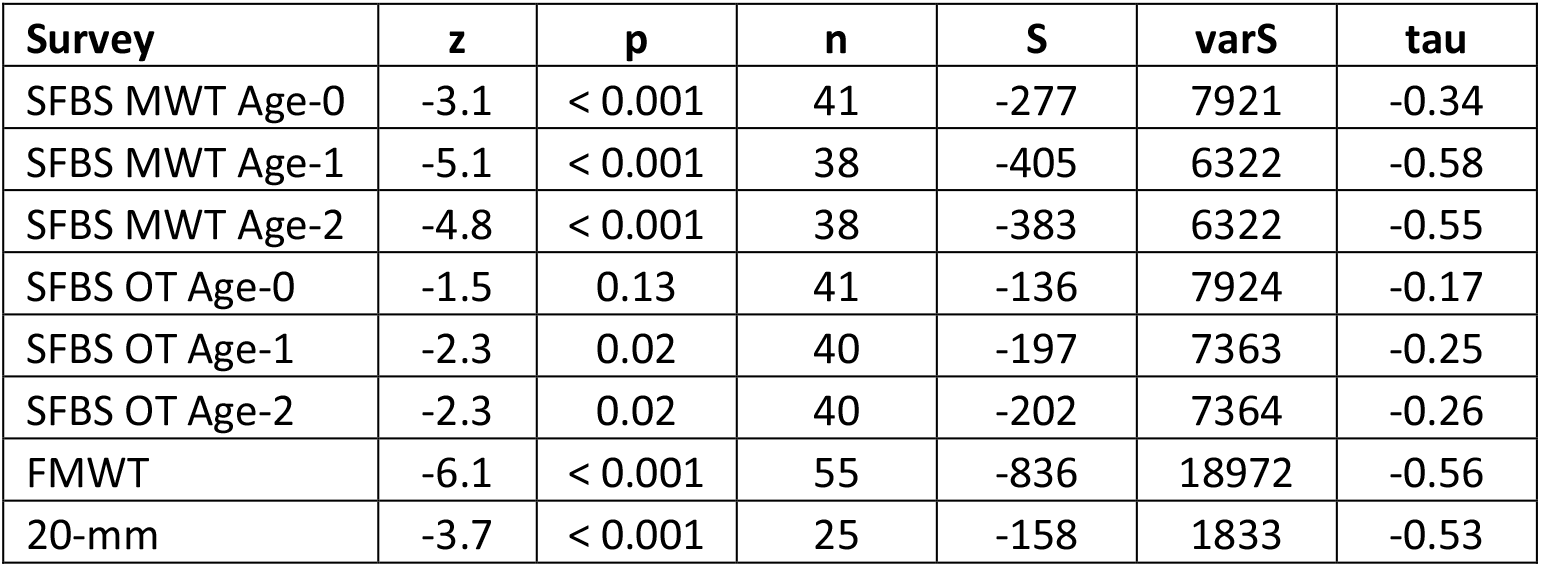
Results of a Mann-Kendall test for monotonic trends in the available time series of abundance indices for Longfin Smelt. Missing values were trimmed from the ends of the time series and missing values within the time series were filled with local average values.

| Survey | z | p | n | S | varS | tau |
| --- | --- | --- | --- | --- | --- | --- |
| SFBS MWT Age-0 | -3.1 | < 0.001 | 41 | -277 | 7921 | -0.34 |
| SFBS MWT Age-1 | -5.1 | < 0.001 | 38 | -405 | 6322 | -0.58 |
| SFBS MWT Age-2 | -4.8 | < 0.001 | 38 | -383 | 6322 | -0.55 |
| SFBS OT Age-0 | -1.5 | 0.13 | 41 | -136 | 7924 | -0.17 |
| SFBS OT Age-1 | -2.3 | 0.02 | 40 | -197 | 7363 | -0.25 |
| SFBS OT Age-2 | -2.3 | 0.02 | 40 | -202 | 7364 | -0.26 |
| FMWT | -6.1 | < 0.001 | 55 | -836 | 18972 | -0.56 |
| 20-mm | -3.7 | < 0.001 | 25 | -158 | 1833 | -0.53 |

### Probability of Extinction over Time

We modified the function extCDF (package popbio; (Stubben and Milligan 2007) to estimate the cumulative density function (CDF) for the probability of extinction. The modified functions use the results of the regression analysis and can incorporate autocorrelation into the simulated abundance at each timestep. The modified function requires the following input values: mean and variance of the population growth rate, the current population size, a quasi-extinction threshold, and a level of autocorrelation. Mean and variance of the population growth rate as well as the quasi-extinction thresholds were set as described above. The value for the current population size was set to the most recent index value that was available (Table 1). A second function produced bootstrapped means and confidence intervals for the CDFs.

To calculate the cumulative probability of extinction over time for the population growth rate derived from the meta-analysis, additional values had to be chosen for the quasi-extinction threshold and the starting population size. We set the initial populations size to the mean of the initial population sizes for the surveys. We set the quasi-extinction threshold to 2.6% of the initial population size to be consistent with the mean ratio of quasi-extinction values and initial population sizes of the surveys, resulting in a quasi-extinction threshold of 50. We considered setting the quasi-extinction threshold to the mean of the quasi-extinction thresholds for all of the surveys, but this value (16) was so low that it risked introducing unaccounted-for positive density-dependent effects (e.g., Allee effects) into the simulation. Realizing that these choices could affect the results of the PVA based on the meta-analysis, we also ran the simulation with several additional values and graphed the results to illustrate the potential effect of these choices.

## Results

### Monotonic Trends

Values for the indices of Longfin Smelt abundance have decreased substantially over the time series (Table 1). None of the abundance indices show evidence of increasing abundance indices: Mann-Kendall tests for all of the abundance indices have negative z-values, indicating negative monotonic trends, and all except for the SFBS age-0 otter trawl index have p-values less than 0.05 (Table 2).

### Population Growth Rates

For all surveys, the most likely direction for annual population growth is negative (Figure 2). Mean population growth rates were less than one for all of the abundance indices, which indicates that population size is declining over time (Table 3); however, variability was high for all surveys and confidence limits on all estimates included the possibility of a stable population size (Figure 2). The mean from the meta-analysis 0.92 (95% CI: 0.86, 0.99), which indicates a population growth rate below that of a stable population. Population growth rates for all surveys suggest that there have been substantial reductions in population size. Based on the mean lambda values, and the assumption that lambda is constant over time, declines over 20 years amount to between roughly 42% and 89% of the population.

**Figure 2:**
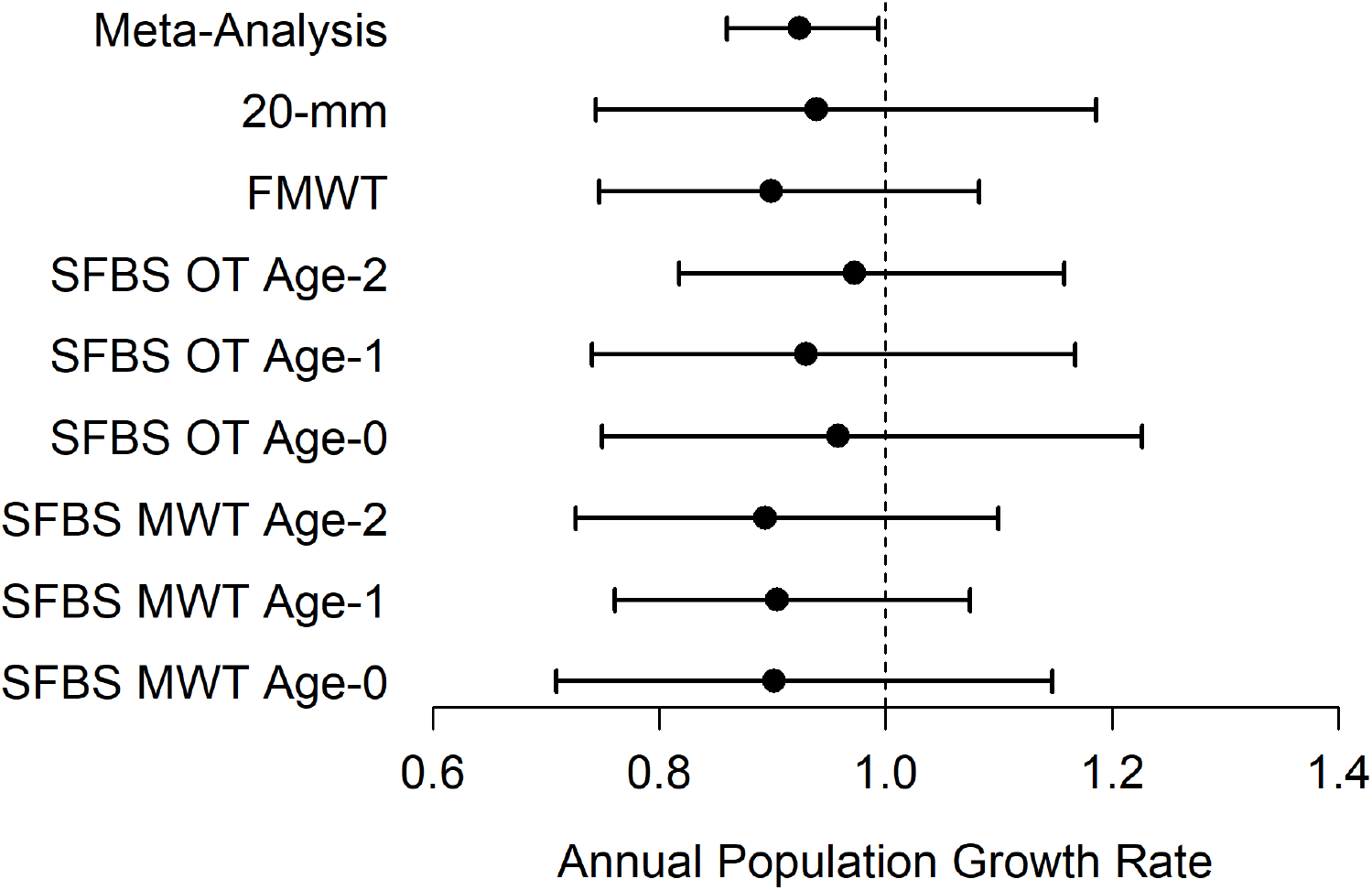
Mean population growth rates from several monitoring programs and a meta-analysis of the mean. Calculations were made using all available years for each survey, based on a count PVA framework. Error bars are 95% confidence limits derived from the regression method developed by Dennis et al. 1991.

**Table 3:**
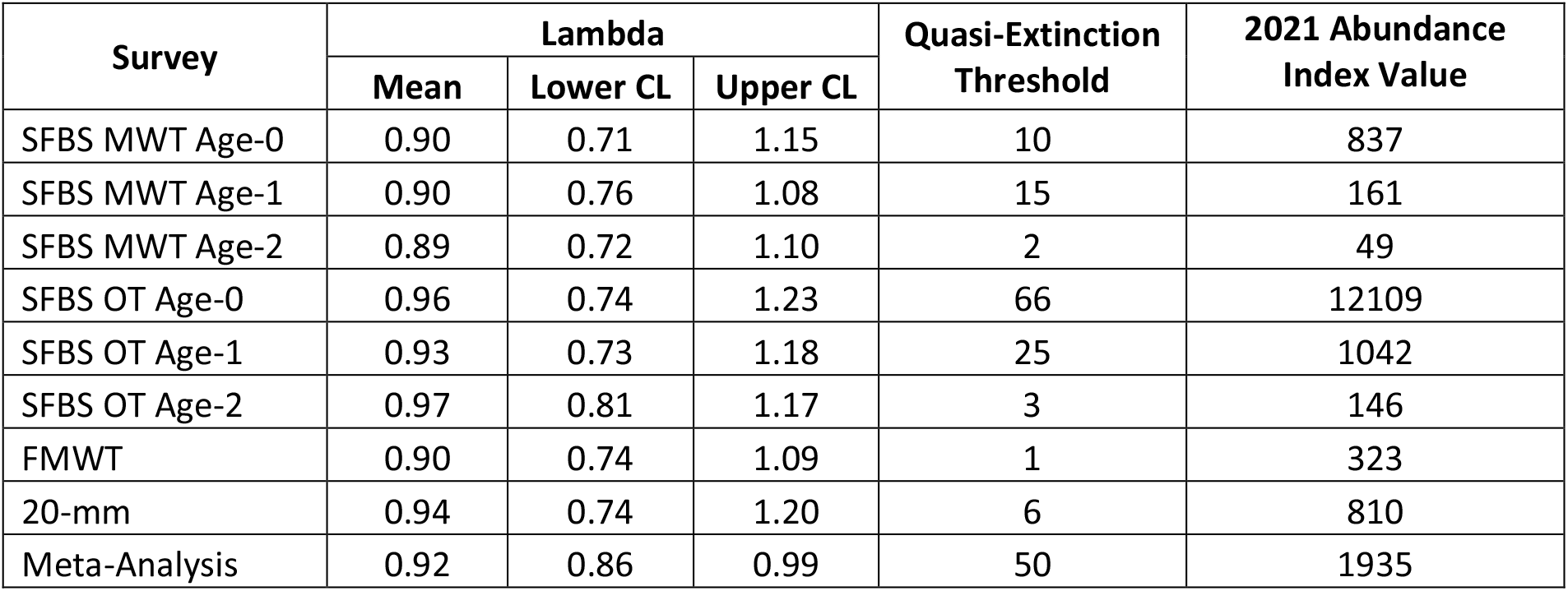
Values corresponding to the mean and 95% confidence limits presented in Figure 1 and the values used to produce Figure 2. These values include all years for which index values were calculated.

| Survey | Lambda |  |  | Quasi-Extinction<br>Threshold | 2021 Abundance<br>Index Value |
| --- | --- | --- | --- | --- | --- |
|  | Mean | Lower CL | Upper CL |  |  |
| SFBS MWT Age-0 | 0.90 | 0.71 | 1.15 | 10 | 837 |
| SFBS MWT Age-1 | 0.90 | 0.76 | 1.08 | 15 | 161 |
| SFBS MWT Age-2 | 0.89 | 0.72 | 1.10 | 2 | 49 |
| SFBS OT Age-0 | 0.96 | 0.74 | 1.23 | 66 | 12109 |
| SFBS OT Age-1 | 0.93 | 0.73 | 1.18 | 25 | 1042 |
| SFBS OT Age-2 | 0.97 | 0.81 | 1.17 | 3 | 146 |
| FMWT | 0.90 | 0.74 | 1.09 | 1 | 323 |
| 20-mm | 0.94 | 0.74 | 1.20 | 6 | 810 |
| Meta-Analysis | 0.92 | 0.86 | 0.99 | 50 | 1935 |

### Probability of Quasi-Extinction over Time

The predicted probability of quasi-extinction for most surveys, except for the 20-mm survey and the SFBS age-2 OT, exceeds 20% by 2040 (Figure 3). Applying the same assumptions over a longer time horizon (i.e., 2050-2065), the suite of surveys predicts that the probability of extinction for the DPS under current conditions is roughly 45-90% across all surveys. Based on the meta-analysis, the mean quasi-extinction value for the population is 33% (25%, 41%) over 20 years and rises to 50% (42%, 58%) in 30 years (Figure 4).

**Figure 3:**
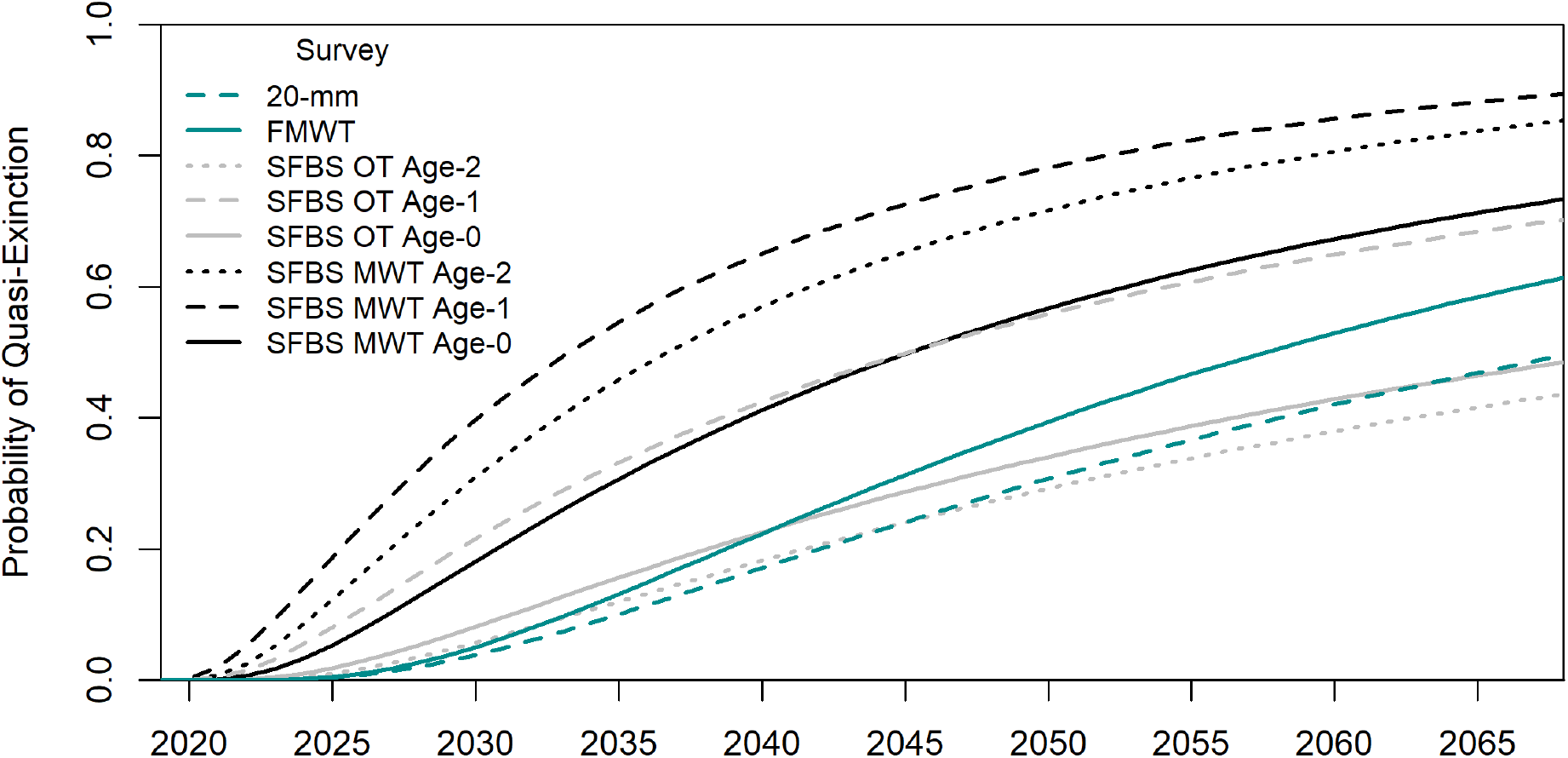
Probability of quasi-extinction for several surveys that report population indices for Longfin Smelt. Values for quasi-extinction thresholds are defined in Table 3. Estimates of population growth rates and variability were derived from all available years for each survey.

**Figure 4:**
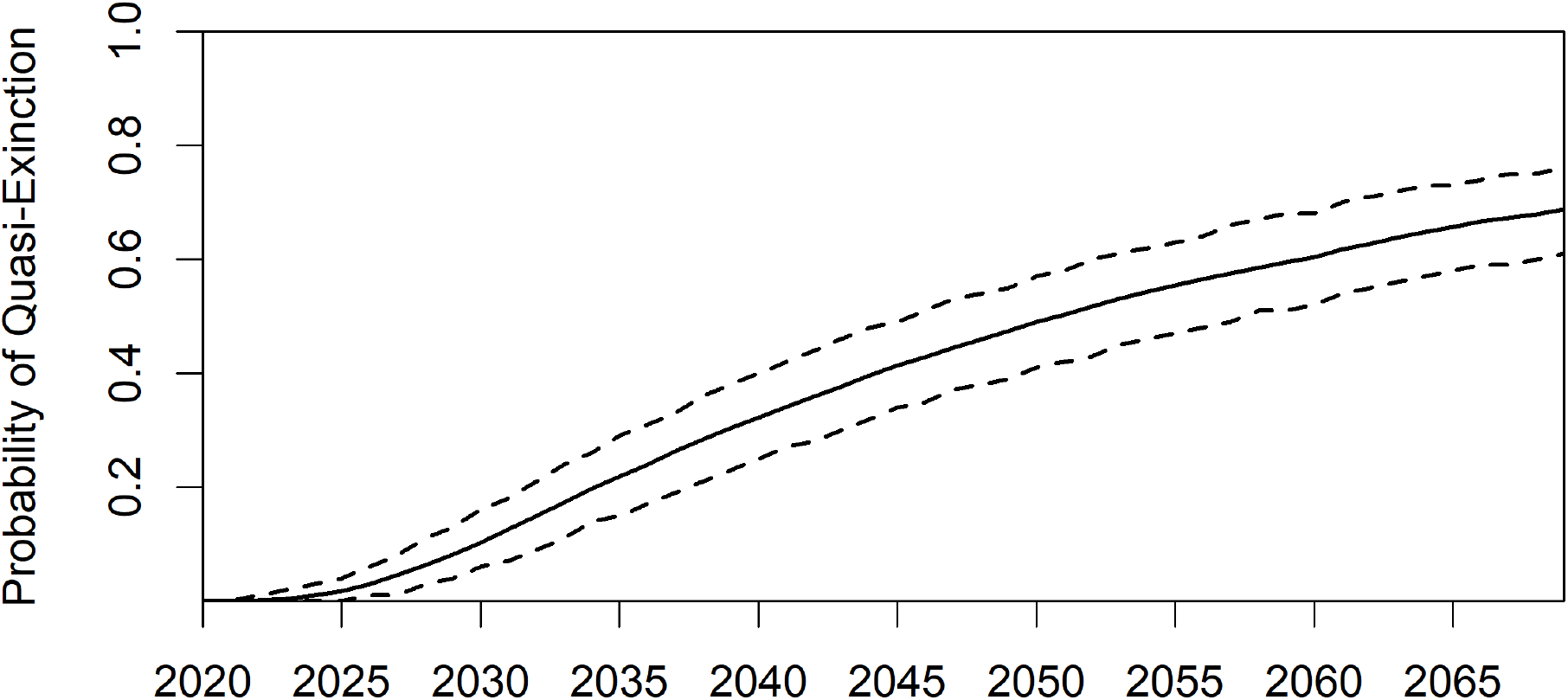
Mean probability of quasi-extinction (solid line), as calculated from the values in the meta-analysis, with bootstrapped 95% confidence bands (dashed lines).

Although the confidence interval on mean population growth rate from the meta-analysis does not include zero, we also calculated the probability of quasi-extinction for a stable population. If the population were not declining (i.e., if the population growth rate were zero), but the population size and variance were the same as calculated for the meta-analysis, the quasi-extinction risk would 40% (31%, 47%) in 20 years and 52% (43%, 59%) in 30 years. Varying the initial abundance could make the extinction probabilities higher, and the values that were tested here reached as high as 80% (Figure 5). The values in Figure 3 were near the lower part of the range of extinction probabilities (high abundance). Quasi-extinction predictions were sensitive to the choice of starting abundance and quasi-extinction threshold (Figure 5). As the starting population size increases, the probability of quasi-extinction rapidly approaches a lower threshold. Increasing the quasi-extinction threshold creates the opposite effect, causing the quasi-extinction probability to reach an upper threshold.

**Figure 5:**
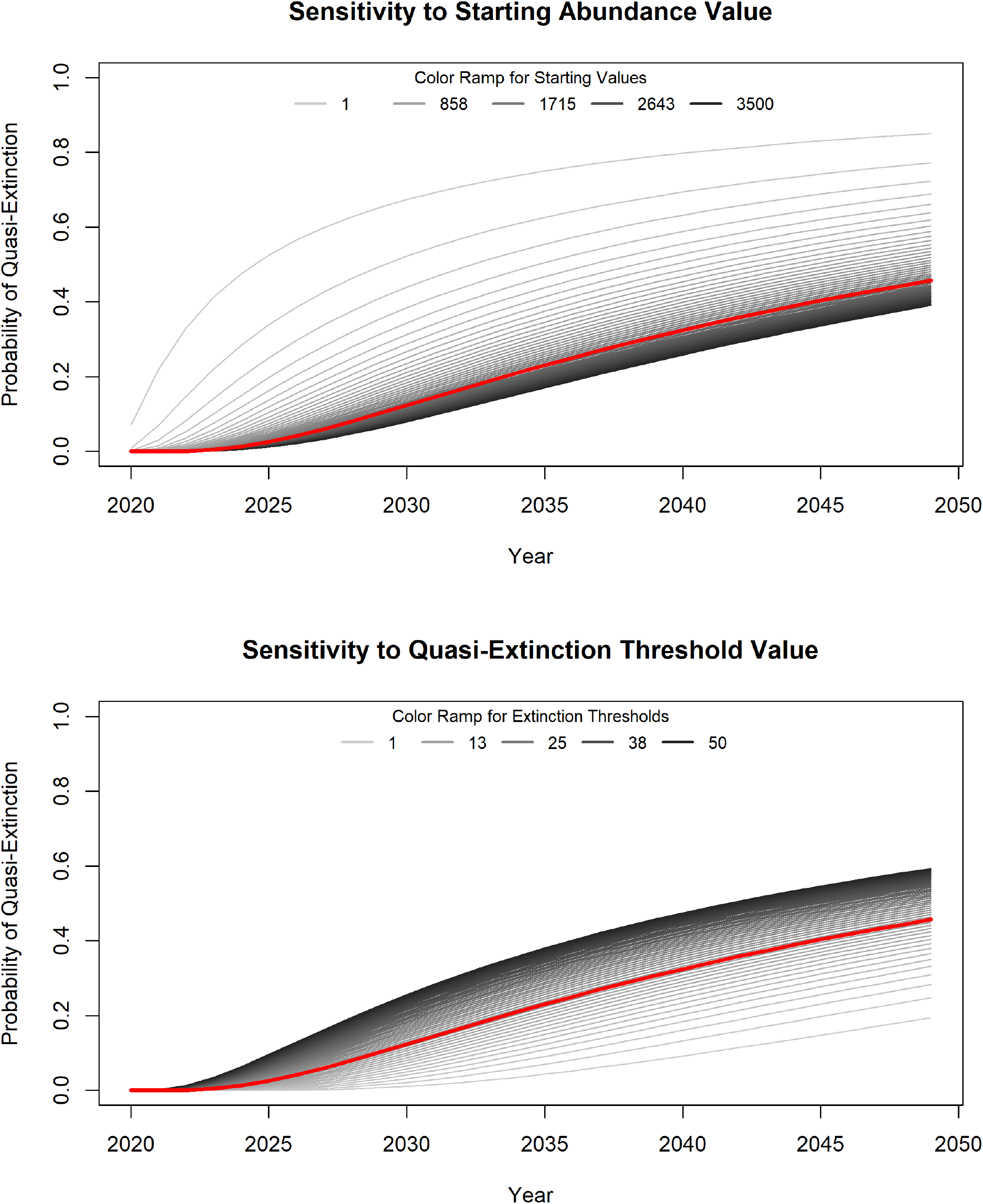
Sensitivity of the probability of quasi-extinction to the choice of quasi-extinction threshold (top) and starting abundance value (bottom). Grayscale lines represent a range of choices. The red line indicates the choice made to produce Figure 3; values for Nc and Ne are given in Table 2.

### Evaluation of Assumptions

Our datasets did not show strong evidence of changes in the mean and variance of population growth rates over time. Plots of population growth rates over time did not show obvious trends in either direction. Similarly, plots of population growth rates as a function of the previous year’s abundance did not show a strong slope, but visual evidence was inconclusive for surveys with outliers. To be sure the linear models were sufficient, we compared them to density-dependent formulations. Model comparisons indicated no evidence to support either of the density-dependent formulations being better than the density-independent formulation. Durbin-Watson tests showed evidence for negative autocorrelation in all indices except the Bay Study age-2 midwater trawl and 20-mm. ACF plots indicated that the strengths of the first-order autocorrelations were about -0.5 when they were significant.

## Discussion

### Population Growth Rates

The suite of long-term monitoring programs in the SFE agrees and provides clear evidence that the abundance on Longfin Smelt in the SFE has declined dramatically over the monitoring record. The abundance index values for recent years are lower than they have ever been and estimates of annual population growth indicate a declining population on average. Longfin Smelt abundance has undergone several step declines since monitoring began (Thomson et al. 2010) as a result of changes in environmental conditions such as the introduction of invasive clams in the mid-1980s (Alpine and Cloern 1992) and the pelagic organism decline (Sommer et al. 2007). The variability in the estimated population growth rates reflects the fact that in some years Longfin Smelt abundance indices increase substantially, but in most years they decrease. Populations with highly variable growth rates tend to have lower levels of population viability over time because the inherent variability tends to make a population grow more slowly over the long term and the population size is more likely to fall below the quasi-extinction threshold than it is for populations with less variable growth rates (Morris and Doak 2002). Without intervention, populations with negative growth rates are expected to go extinct, regardless of the initial population size or the variability in their growth rates. The main question to investigate becomes when extinction is likely to happen, because this determines the timeframe for implementing management actions to increase the growth rate.

Information from multiple surveys can help us be more confident in our summaries (and thus our decisions). The meta-analysis indicates that on average, the surveys are tracking an actual decrease in Longfin Smelt abundance over time confirming the apparent visible data trend. This finding is consistent with previous studies that used indices of abundance to evaluate patterns in Longfin Smelt populations. For example, a pattern of decreasing catch per trawl has been fairly consistently over time, with a few short increases (Sommer et al. 2007; Thomson et al. 2010). By leveraging information from multiple monitoring surveys, we obtain better information in the form of more precise estimate of the population growth rate. When using information from all of the surveys together, the variability around the mean is dampened, compared to the individual surveys, and the confidence bands no longer include zero. Variation around the population growth rate contributes to high estimated risk of quasi-extinction, whether predictions are made using the mean population growth rate or using an assumption that population growth is stable. In either scenario, the message for managers is that increased population growth rates and reducing variation around those growth rates are necessary to reduce the risk of quasi-extinction.

### Evaluation of assumptions

PVAs are best used to predict extinction probabilities when extensive and reliable data exist, and if the distribution of vital rates is stationary or predictable in the future (Coulson et al. 2001). For Longfin Smelt, several long-running datasets exist that can inform a PVA. Although the structure of some PVAs can be very complex to account for various issues with data quality or for a lack of data, here we took a simple approach, in part because we have an abundance of consistent data. Our approach leverages the same assumptions as are used for interpreting monitoring data in general, specifically that indices of abundance are proportional to true abundance. This assumption is necessary otherwise monitoring data are not useful. Because of this, this analysis may be more applicable to evaluating the surveys themselves than the abundance of Longfin Smelt directly. Future work will address this by leveraging on-going efforts to calculate absolute abundance for Longfin Smelt. Another assumption for PVAs is that there is no observation error in the time series. The EGSS model accounts for observation error that is present in the timeseries for the surveys. Removing observation error guards against introducing unrealistic variability in the estimates of the population growth rate, which would lead to underestimating viability and thus pessimistic predictions (Staples et al. 2004; Humbert et al. 2009). The meta-analysis approach further dampens this effect because incorporating information from all of the surveys to improves the precision of the estimate of the population growth rate overall.

Different assumptions about the form of population regulation (i.e., density dependence) have been shown to affect the expected population viability significantly (LaMontagne et al. 2002). Our analysis relies on a simplified version of the population dynamics for Longfin Smelt. In particular, it does not account for age structure in the population or potential density dependence in the time series of recruits per spawner, which has been suggested by previous studies (Nobriga and Rosenfield 2016). Our evaluation of the data supports a density-independent modeling structure when examining year to year variation of the same age class. Recent abundance levels are very low so it is unlikely that negative density dependence would affect our forecasted simulations, especially those using the meta-analysis mean and SD values. Even if high population values were predicted, the effect of negative density dependence would not change our conclusions that Longfin Smelt are at risk for extinction. Left unaccounted for, negative density dependence could cause unrealistically high estimates of abundance, which would make the calculated quasi extinction rates lower than they should be (i.e., the population would appear less likely to go extinct than it should). Longfin Smelt might already be experiencing Allee effects (i.e. positive density dependence). The selected quasi-extinction thresholds were chosen to represent values that are much lower than recent averages and to be tailored to the magnitudes of individual indices, but they were not informed by considerations of when management actions could still be effective for managing population sizes. As a result, quasi-extinction thresholds should be examined carefully with management actions in mind during any potential management planning.

### Management Implications

The ESA does not set quantitative definitions or thresholds for applying threatened or endangered designations, but other benchmarks can provide context for interpreting quasi-extinction rates. For example, a 20% risk of extinction over 20 years has been proposed by multiple sources as a criterion for a species being endangered (Lindley et al. 2007; Mace et al. 2008). Another definition holds that a population with a 95% probability of persisting for 100 years can be considered secure and not at risk of extinction (Hamilton and Moller 1995). We found that the Longfin Smelt population in the San Francisco Estuary meets these criteria to be considered at risk of extinction. For Longfin Smelt, most of the increase in cumulative probability of extinction occurs in the next two decades, but there may be time to make changes before Longfin Smelt become undetectable by long-term monitoring surveys. In cases like that of Longfin Smelt where the population size is small, but the risk of extinction is relatively small for the immediate future, conservation and management efforts should focus on achieving long-term viability by taking steps to increase the population growth rate for the species (Morris and Doak 2002). The structure in the population growth rates can also suggest some management strategies. Negative autocorrelation in the time series of population growth rates indicates that Longfin Smelt populations tend to decrease after good years, thus recovery from the current downward trajectory will require conditions that produce a series of good years in a row.

This study demonstrates a practical way that having multiple sources of information creates better information about the trajectory of the population. Individually, the monitoring surveys contribute information about specific life stages or ages to our understanding of the population. Here we demonstrated that, as a data synthesis, combining information from multiple surveys improves our confidence in our estimate of the trajectory of the population. Scientists who produce and track the information provided by several indices of abundance develop an intuition about the trajectory of the species. The methods we used here provide a quantitative means to summarize the process that creates that intuition. In effect, we show how a set of simple PVAs used as preliminary models can formalize uncertainty in our current understanding of the trajectory of the species (Hamilton and Moller 1995).

Even when multiple data sources identify likely species declines, synthesizing the landscape of information into one metric and an associated graphical summary may be useful for ecosystem managers and decision-makers. In addition to succinctly communicating risk information, this summary presents a benchmark for evaluating future decisions. This PVA represents a baseline scenario that considers a future with no changes in management or environmental conditions, which can be used for comparison with potential future management actions and recovery planning. One limitation of the current study for management implications is the wide confidence intervals, even on the meta-analysis mean. We addressed this issue by bootstrapping estimates, but it could also be addressed through more sophisticated models that account for variability in a more structured way. A Bayesian state-space model approach to combining data from multiple surveys has been shown to produce precise parameter estimates, even when data were sparse (e.g., Johnson et al. 2010) and we plan to pursue this approach in an upcoming manuscript.

## Acknowledgements

The authors thank the many members of the California Department of Fish and Wildlife’s Interagency Ecological Program monitoring teams, both current and former, who collected the data and calculated the indices that we used in this manuscript. We also thank the many people who improved early versions of this manuscript through the review processes associated with the development of the species status assessment and the proposed rule for the Bay-Delta Distinct Population Segment of Longfin Smelt. Comments from internal reviewers, those who participated in peer and partner review, and those who submitted reviews through the public comment period helped shape this paper. Funding for this work was provided by the authors’ respective agencies. The findings and conclusions in this article are those of the authors and do not necessarily represent the views of the U.S. Fish and Wildlife Service or the California Department of Fish and Wildlife.

## Supplemental Information

**Table S1:** Model comparisons for different density-dependence formulations.

| Survey | Model | df | AICc | Converged | r | R_p | K | K_p |
| --- | --- | --- | --- | --- | --- | --- | --- | --- |
| FMWT | Beverton-Holt | 3 | 212.12 | TRUE | 0.14 | 0.42 | -680.10 | 0.40 |
| FMWT | Ricker | 3 | 213.03 | TRUE | -0.21 | 0.47 | 8597.13 | 0.44 |
| FMWT | Linear | 3 | 213.03 | TRUE | 0.00 | 0.14 | NA | NA |
| mwt.0 | Beverton-Holt | 3 | 160.15 | FALSE | 1003.09 | 1.00 | 86.04 | 0.85 |
| mwt.0 | Ricker | 3 | 159.17 | TRUE | -0.19 | 0.64 | 42811.97 | 0.62 |
| mwt.0 | Linear | 3 | 159.17 | TRUE | 0.00 | 0.22 | NA | NA |
| mwt.1 | Beverton-Holt | 3 | 122.52 | FALSE | -286790.80 | 1.00 | -74.62 | 0.89 |
| mwt.1 | Ricker | 3 | 121.43 | TRUE | -0.27 | 0.31 | 52207.57 | 0.36 |
| mwt.1 | Linear | 3 | 121.43 | TRUE | 0.00 | 0.29 | NA | NA |
| mwt.2 | Beverton-Holt | 3 | 101.12 | TRUE | 0.83 | 0.43 | 53.10 | 0.64 |
| mwt.2 | Ricker | 3 | 102.37 | TRUE | 0.08 | 0.78 | 1010.97 | 0.70 |
| mwt.2 | Linear | 3 | 102.37 | TRUE | 0.00 | 0.63 | NA | NA |
| ot.0 | Beverton-Holt | 3 | 159.76 | FALSE | 385.39 | 1.00 | 66.88 | 0.93 |
| ot.0 | Ricker | 3 | 159.14 | TRUE | -0.09 | 0.78 | 17670.29 | 0.76 |
| ot.0 | Linear | 3 | 159.14 | TRUE | 0.00 | 0.39 | NA | NA |
| ot.1 | Beverton-Holt | 3 | 147.24 | TRUE | 0.19 | 0.47 | -4780.19 | 0.42 |
| ot.1 | Ricker | 3 | 148.16 | FALSE | NA | NA | NA | NA |
| ot.1 | Linear | 3 | 148.10 | TRUE | 0.00 | 0.91 | NA | NA |
| ot.2 | Beverton-Holt | 3 | 115.15 | TRUE | 0.02 | 0.90 | -138.66 | 0.89 |
| ot.2 | Ricker | 3 | 116.97 | FALSE | NA | NA | NA | NA |
| ot.2 | Linear | 3 | 115.75 | TRUE | 0.00 | 0.39 | NA | NA |
| 20-mm | Beverton-Holt | 3 | 65.63 | FALSE | 1354.41 | 1.00 | 25.26 | 0.81 |
| 20-mm | Ricker | 3 | 67.87 | FALSE | NA | NA | NA | NA |
| 20-mm | Linear | 3 | 65.32 | TRUE | 0.00 | 0.31 | NA | NA |

**Figure S1:**
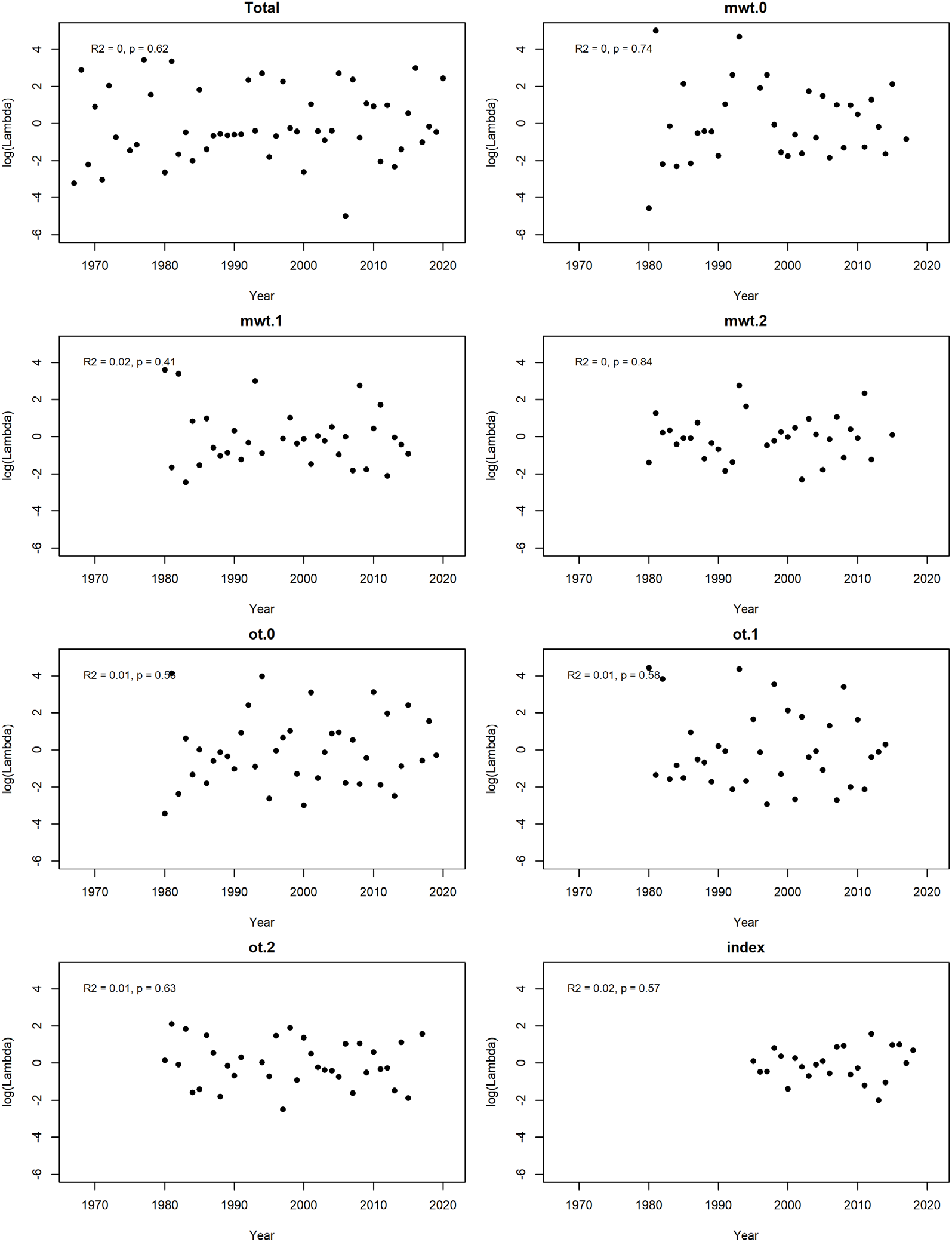
Graphs and regression statistics for checking the assumption that population growth rates to not change over time.

**Figure S2:**
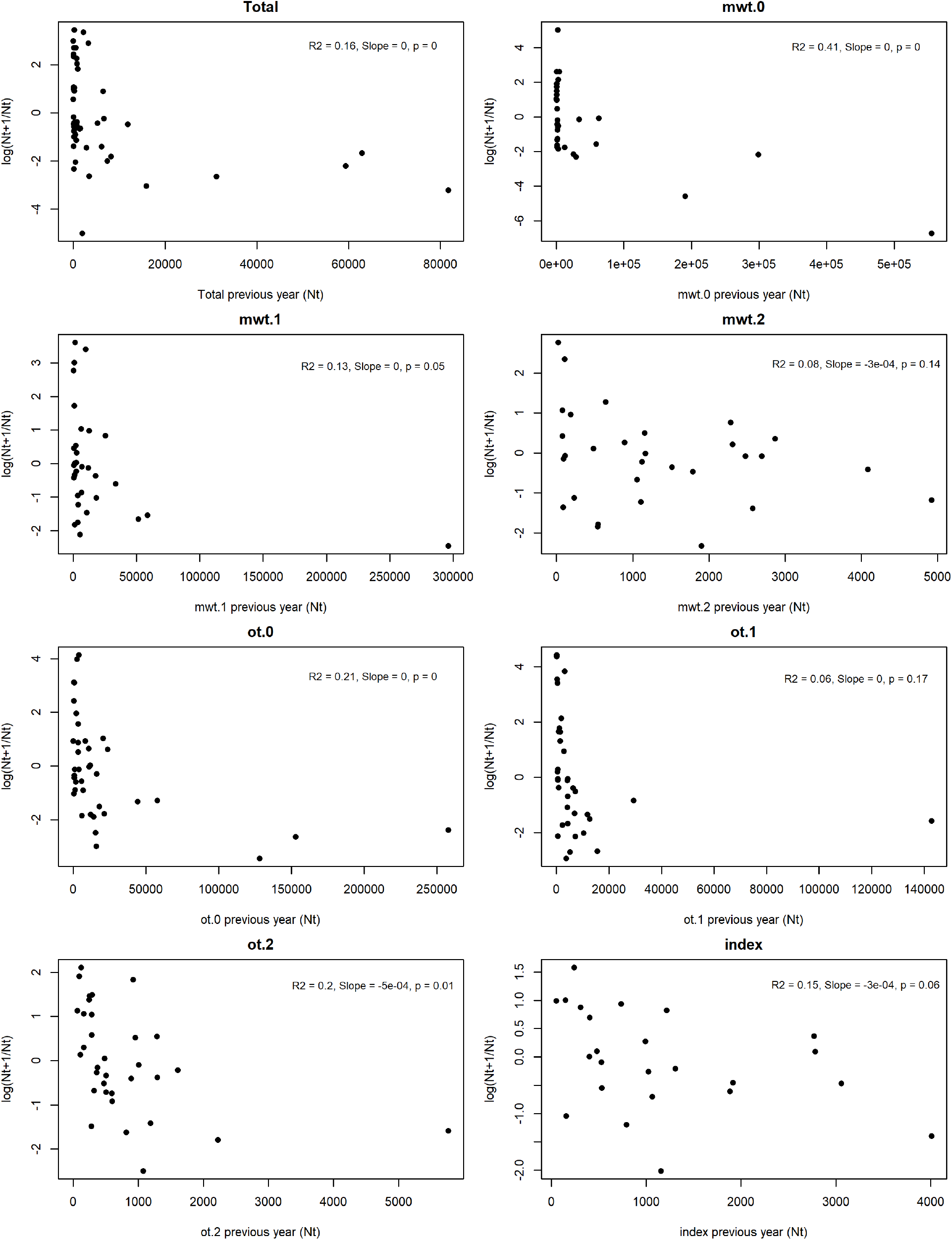
Graphs and regression statistics for checking whether the population growth rates are density dependent.

**Figure S3:**
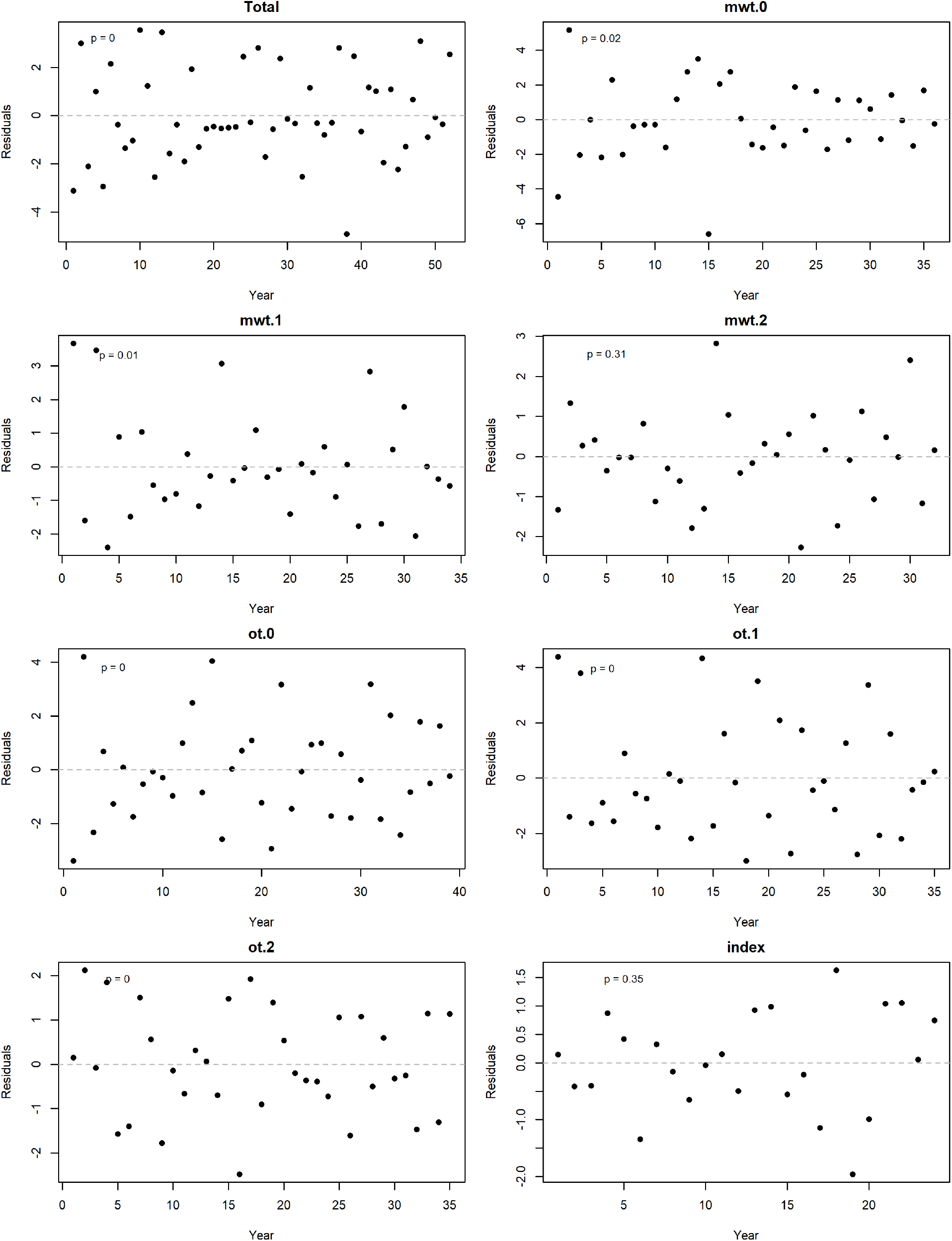
Graphs and statistics for checking for environmental autocorrelation. Results of Durbin-Watson tests for first-order autocorrelation are represented as p-values printed on each panel.

